# Universal scaling across biochemical networks on Earth

**DOI:** 10.1101/212118

**Authors:** Hyunju Kim, Harrison B. Smith, Cole Mathis, Jason Raymond, Sara I. Walker

## Abstract

The application of network science to biology has advanced our understanding of the metabolism of individual organisms and the organization of ecosystems but has scarcely been applied to life at a planetary scale. To characterize planetary-scale biochemistry, we constructed biochemical networks using a global database of 28,146 annotated genomes and metagenomes, and 8,658 cataloged biochemical reactions. We uncover scaling laws governing biochemical diversity and network structure shared across levels of organization from individuals to ecosystems, to the biosphere as a whole. Comparing real biochemical networks to random chemical networks reveals the observed biological scaling is not solely a product of the biochemistry shared across life on Earth. Instead, it emerges due to how the global inventory of biochemical reactions is partitioned into individuals. We show the three domains of life are topologically distinguishable, with > 80% accuracy in predicting evolutionary domain based on biochemical network size and average topology. Taken together our results point to a deeper level of organization in biochemical networks than what has been understood so far.

---

There is increasing interest in whether biology is governed by general principles, not tied to its specific chemical instantiation or contingent upon evolutionary history ^1–3^. Such principles would be strong candidates for being universal to all life ^4,5^. Universal biology, if it exists, would have important implications for our search for life beyond Earth ^6–9^, for engineering synthetic life in the lab ^10,11^, and for solving the origin of life ^12,13^. Systems biology provides promising quantitative tools for uncovering such general principles ^14–16^. So far, systems approaches have primarily focused on specific levels of organization within biological hierarchies, such as individual organisms ^17,18^ or ecological communities ^19,20^, and are rarely applied to the biosphere as a whole. But, biology exhibits some of its most striking regularities moving up in levels of organization from individuals to ecosystems, and these regularities may only truly manifest at the level of ecosystems, and ultimately the biosphere ^21,22^. For example, while individual organismal lineages fluctuate through time and space, the functional and metabolic composition of ecological communities is dynamically stable ^23,24^. To understand the general principles governing biology, we must understand how living systems organize across levels, not just within a given level ^25,26^.

In order to explore regularities within and between levels of organization, we adopt a network view of biochemistry ^17,27–29^ by constructing biochemical reaction networks from genomic and metagenomic data. We show biochemical networks share universal organizational properties across levels, characterized by scaling laws determining how topology and biochemical diversity change with network size. These scaling relations exist *independent* of evolutionary domain or level of organization, applying across the nested hierarchy of individuals, ecosystems, and the biosphere. The biochemical diversity and network properties driving this scaling behavior are predictive of evolutionary domain, indicating the biochemical network structure for each domain is distinct even though all three conform to the same scaling behavior. Our results provide a first quantitative demonstration the application of network theory at a planetary scale can uncover properties existing across different levels of organization within the biosphere, and can be predictive of major divisions within a given level (such as domains). Taken together these results provide new paths forward for identifying universal properties of life.

Our analysis begins with a global database of genomes and metagenomes, sampled from across life on Earth. We leverage available existing annotated genomic data representing the three domains of life, including genomes of 21,637 bacterial taxa and 845 archaeal taxa from the Pathosystems Resource Integration Center (PATRIC) ^30^, and 77 eukaryotic taxa from the Joint Genome Institute (JGI) ^31^. Our metagenomic data includes 5,587 metagenomes from JGI cataloging ecosystem-level biochemical diversity across the planet, see Fig. 1.

**Figure 1:**
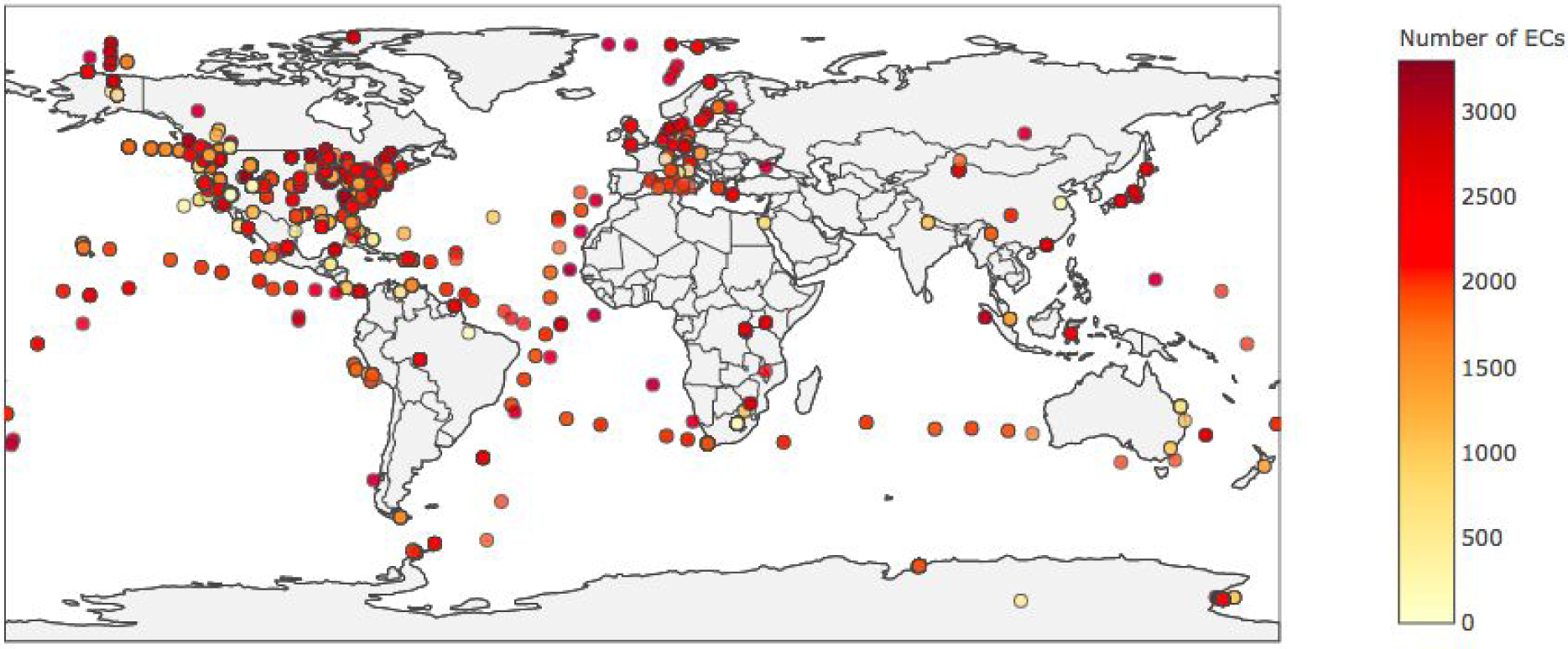
Enzyme diversity of ecosystems across Earth. Shown is the geographical distribution of 5,587 ecosystems, colored by the number of different enzyme functional classes (enzyme commission (EC) numbers) encoded in sampled metagenomes (from JGI). Despite large variances in the enzyme diversity and what enzymes are present in each ecosystem, all ecosystems sampled are found to conform to the same scaling behavior for biochemical diversity and topology as a function of biochemical network size, see Fig. 3.

From this data, we constructed biochemical networks for each individual organism (genome) and ecosystem (metagenome) using reaction data cataloged in the Kyoto Encyclopedia of Genes and Genomes (KEGG) ^32^. Building on prior work studying biosphere-level models of metabolism ^33–35^, we use the database of all 8,658 enzymatically catalyzed reactions cataloged in KEGG as a proxy for the biochemistry of the biosphere as a whole, modeled as a ‘soup of enzymes’ by disregarding the boundaries of individual species^19^. Network representations of ecosystem-level and biosphere-level biochemistry are ‘compartment-free’ in that no knowledge of individual species is included. Previous topological analyses of biochemical networks have primarily focused on the subset of biochemical reactions associated with metabolism ^28,29^. Since we are interested in properties universal across life, and not just subsets of living processes, we instead construct networks inclusive of every known catalyzed reaction (regardless of pathway) coded by the respective genome or metagenome, provided the reaction is cataloged in KEGG.

Adopting a network representation allows systematic quantification of topological properties using graph theory and statistical mechanics^16,17,36–41^. Using two different graph projections, we compare biochemical networks to test whether they are similar or different across levels (see Methods for details on network construction). A widely implemented framework for assessing commonality across different systems is to look at their scaling behavior ^42–47^. If scale-invariant properties are found, it can be suggestive of deeper, underlying organizing principles ^3,48,49^, such as when power-law scaling emerges at criticality in thermodynamic phase transitions ^50^. We therefore sought to determine whether biochemical networks display similar scaling laws governing their topology and chemical diversity across levels, indicative of the existence of self-organizing principles universal across different biological levels.

There are three alternative scenarios to be tested relating network structure across individuals, ecosystems and the entire biosphere, each is shown in Fig. 2. In the first, biochemistry does not have shared network structure across levels, and different scaling behaviors emerge at different levels. In the second scenario, biochemistry has shared network structure across levels, but this shared structure can be fully explained by the structure of random chemical networks (generated from random collections of chemical reactions used by biology). In this case, biochemical networks would be statistically indistinguishable from random chemical networks, implying the self-organizing principles are solely chemical and not biological in origin. In a third scenario, biochemistry has shared structure across levels, which is different from that of random chemical networks. We find the third scenario to be consistent with our analysis, suggesting the presence of universal organizing principles unique to biology that recur across biological levels of organization.

**Figure 2:**
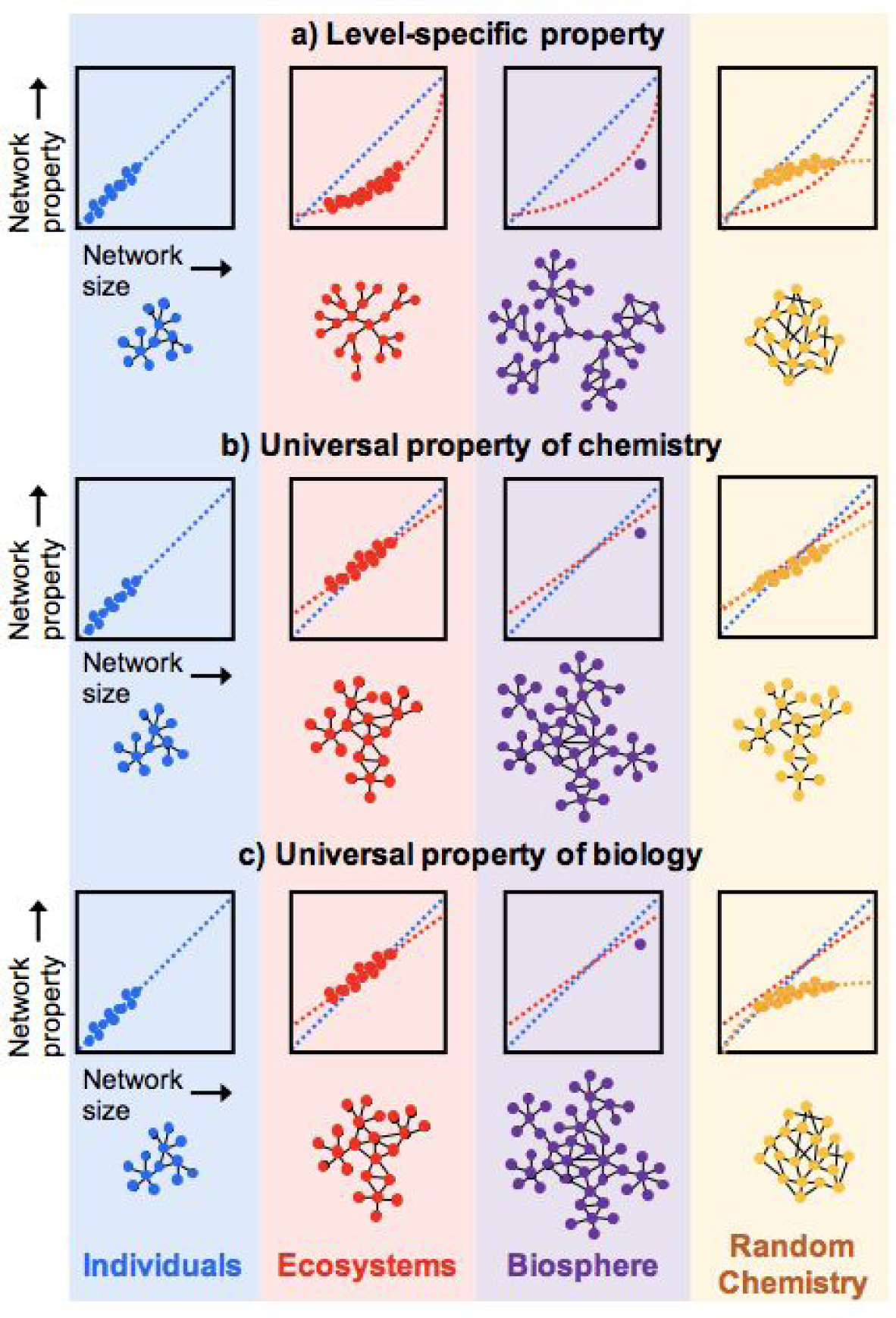
Three alternative scenarios for how biochemical network structure might be similar or dissimilar across levels of organization. For each scenario, illustrative plots show examples of scaling behavior of a network property as function of network size, where each data point corresponds to the measure for a single instance of a network. In the first (a) biochemistry does not exhibit common network structure across levels, and different properties emerge at different levels. In the second (b), biochemistry has a common network structure across all levels, but this structure is also shared by random chemical networks. In the final scenario (c), biochemistry has shared structure across all levels, which is different from that of random chemical networks. Our results are consistent with this third scenario, indicative of universal organizing principles recurring across biological levels, which are unique to biology (not shared by random chemistry).

Before proceeding to the details of our results, it is worth noting the well known challenges associated with the introduction of statistical artifacts when coarse-graining real-world systems to generate graphical representations ^51,52^. For example, bipartite network representations of biochemistry (treating reactions and substrates as two disjoint sets of nodes) have information that cannot be recovered from unipartite representations (which treat only substrates as nodes). The challenge of choosing a projection arises because biochemical networks are themselves a multi-layer system consisting of enzymes and their catalyzed reactions; enzymes (often abstracted away in network representations) control the biological organization we aim to characterize. To ensure the regularities we report are reflective of the true underlying organization of biochemistry, and are not statistical artifacts introduced by a specific choice in coarse-graining, we therefore consider both a unipartite and bipartite projection in our analysis ^51^. We also compare catalytic diversity - quantified in terms of the number of enzymes and reactions - across levels, which are independent of network representation. As we will show, common scaling laws describing biochemical networks across levels of organization are consistently observed in each of these different views of biochemistry, confirming our results are independent of the type of network representation.

One remaining consideration once a network representation is adopted is how to analyze it. So far, the majority of network analyses applied to biochemistry have focused on the ‘scale-free’ structure of metabolic networks^27,53,54^. For example in a seminal paper by Jeong et al. ^27^, it was shown (using a unipartite representation) that metabolic networks from all three domains of life exhibit the characteristic power-law degree distribution of a scale-free network, with similar scaling exponents for bacteria, archaea and eukaryotes. This and other previous work has focused primarily on properties ***within*** single instances of a network (e.g. an individual organism’s metabolism). However, as we stated earlier, our interest is in looking at properties ***across*** networks (e.g. describing biochemical networks at the individual and ecosystem-level). We therefore focus on topological measures such as average shortest path length, average clustering coefficient and assortativity (degree correlation coefficient), which are directly comparable across different networks, allowing us to make statements about regularities existing across biochemistry sampled from different levels of organization.

## Results

### Shared scaling laws describe biochemical networks and catalytic diversity across levels of organization

Organisms can vary widely in their number of genetically encoded reactions, and ecosystems generally include more encoded reactions than individuals. We therefore compare topological properties relative to the size of biochemical networks as a relevant scaling parameter for our analysis. We define network size as the number of molecules connected through catalyzed reactions within the largest connected component (LCC) for a given biochemical network. We focus analyses on the LCC since some measures are not defined on disconnected networks. The LCC includes > 90% of compounds for all but the smallest networks in our study, and >97% of compounds for the largest (see Methods on *Topological measures*, Supplementary Fig. S1, and Supplementary Table S1). The fact that the LCC is not 100% of the network could be attributable to missing data in the annotation of genomes and metagenomes. We therefore verified our results are not sensitive to similar magnitude of missing data by confirming the scaling trends reported here are not affected when 10% of nodes are randomly removed (see Supplementary Fig. S2).

We calculate several frequently implemented topological measures for the LCC for each network. We classify properties (e.g. topological or diversity measures) as *universal* when they scale in the same way across levels. We identify these cases by properties which scale according to the same fit across levels (e.g. network average clustering coefficient scales linearly with network size for both individuals and ecosystems, and we thus identify this scaling as universal across levels). Shared fit functions across levels suggest mechanisms driving the structure of biochemical networks may be independent of level of organization; in such a case individuals and ecosystems could both be subject to the same general principles acting to architect them. That is, we do not require the scaling coefficients to be exactly the same (indicating the tuning of mechanisms generating structure in individuals and ecosystems), but we do require the same fit to be shared across our data (indicating the possibility of shared generative mechanisms) to qualify as universal.

To test whether biology exhibits universal scaling behavior across levels, we first determined how topological properties and biochemical diversity vary with size for all individuals and ecosystems in our data set. Measured values for the unipartite representation and catalytic diversity (enzymes, reactions) are shown in Fig. 3 as a function of network size (see Supplementary Fig. S4 for data on bipartite representation which exhibits similar consistency across levels). We find individuals and ecosystems scale according to the same functional form for each network and diversity measure, with similar scaling coefficients (for fits and confidence intervals, see Supplementary Data S1). Scaling for individuals and ecosystems is therefore universal. For some measures (assortativity and betweenness) the biosphere falls within the 95% confidence interval observed for fits of ecosystem level scaling. An exception is clustering coefficient, where the biosphere significantly departs from the observed ecosystem scaling: this could be be attributable to missing data on global enzymatic diversity (which falls slightly below what our scaling laws would predict). Topological measures that scale following power-law fits (y = y_0_ x^β^, where β=βind for individuals and β=β_eco_ for ecosystems) include: average betweenness (βind=-1.1581, β_eco_=1.136), average shortest path length (β_ind_=-0.117, β_eco_=-0.084), and number of edges (βind=1.219, β_eco_=1.243). Both biochemical diversity measures also scale according to power-law fits: number of enzyme classes (a proxy for enzymatic diversity) (β_ind_=1.294, β_eco_=1.838), and number of reactions (β_ind_=1.229, β_eco_=1.319). Average clustering coefficient scales with a linear fit (y = mx + y0, m=mind for individuals and m=m_eco_ for ecosystems) for individuals and ecosystems (mind=3.77×10^−5^, meco=3.32×10^−5^). These results rule out the possibility scaling laws are level-specific (Fig. 2, row A). The observed scaling laws confirm biochemical networks exhibit shared structure across levels of organization, where network properties and diversity are largely determined by size as the relevant scaling parameter.

**Figure 3:**
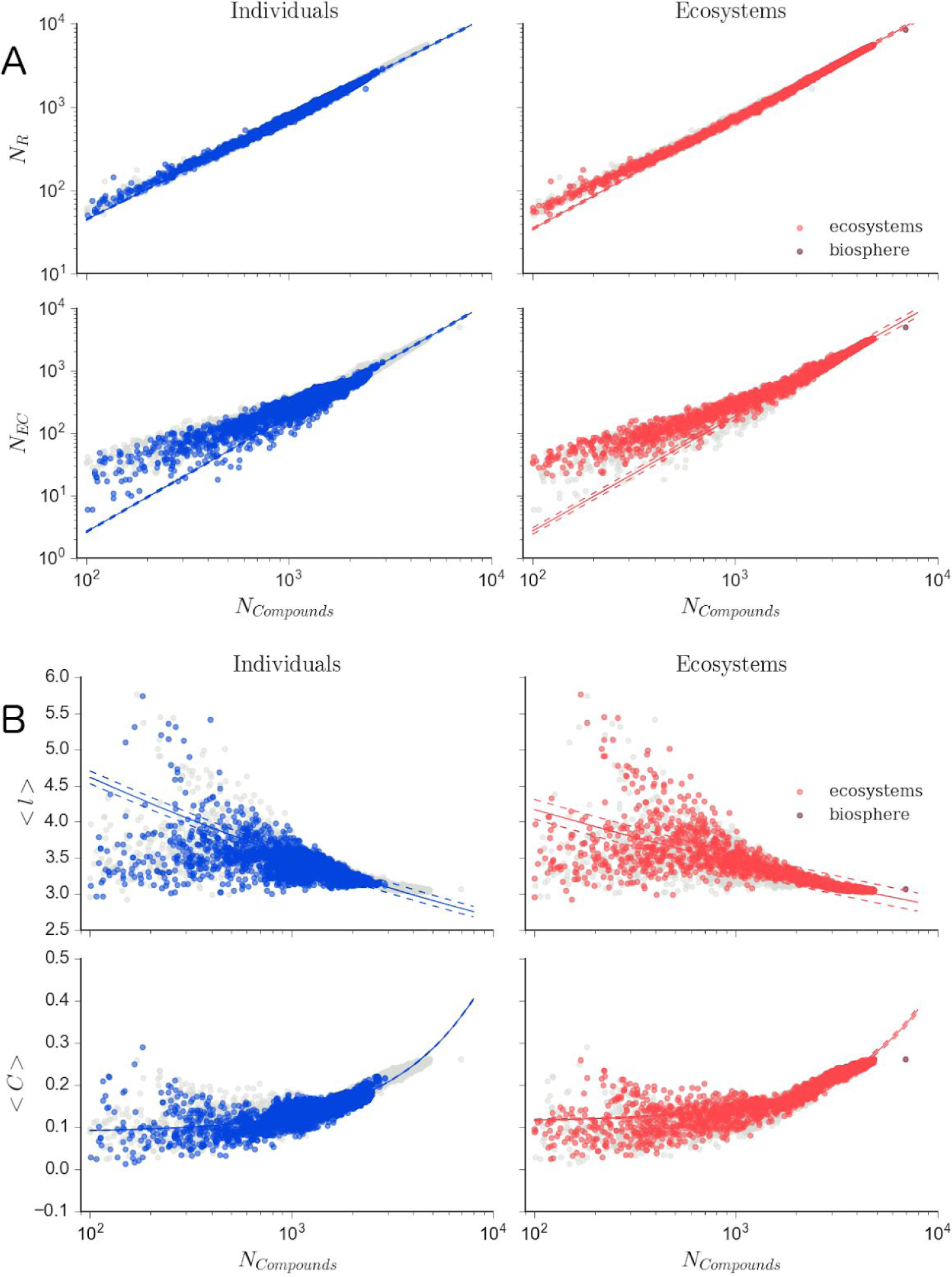
Common scaling laws describe biochemical networks across levels of organization. Scaling of biochemical measures for individuals (left column) and ecosystems (right column) shares the same functional form for catalytic properties (such as enzyme and reaction diversity) and for topological measures. (A) Shown from top to bottom are number of reactions (*N*_*R*_), and number of enzyme classes (*N*_*EC*_). (B) Shown from top to bottom are average shortest path (*<l>*), and average clustering coefficient (*<C>*). All measures are as a function of the size of the largest connected component (*N*_*Compounds*_). Ecosystems include metagenomes (red) and the biosphere-level network (maroon). Fits for each dataset (solid lines) are shown with 95% confidence intervals (dashed lines). For reference, shown in light grey is data for all biochemical networks (individuals, ecosystems, biosphere). Additional measures are shown in Supplementary Fig. S3, and scaling for bipartite networks is shown in Supplementary Fig. S4.

### Real networks exhibit different scaling behavior than random chemical networks

The observed universal scaling across individuals and ecosystems could be unique to biology, or it could arise due to self-organizing principles of chemistry. If the later is true, we should expect random chemical networks to exhibit the same fit functions as real biochemical networks do. Testing this requires comparison to random chemical networks, which much be generated with an appropriate control to be informative ^55^. Since we are interested in the global organization of biochemistry, we constructed control random chemical reaction networks (henceforth called random reaction networks) by merging randomly sampled reactions from the KEGG database (see Methods for details on network construction). This random sampling produces ensembles of random chemical networks which globally (over the ensemble) share the same chemistry as our biosphere. These networks are composed of biochemical reactions, but with no notion of individuals as ‘units’ of selection. Most highly connected nodes (participating in many reactions) are common to all three domains, e.g. ATP and H_2_O ^27,56^. Therefore, this uniform sampling procedure yields random control networks that tend to include the most common compounds used by life.

We performed the same analyses on the random reaction networks as real biochemical networks. We observe random reaction networks do not scale according to the same functional form as biochemical networks for many network and diversity measures (Fig. 4). The fits for average clustering coefficient of random reaction networks favor a power-law function, compared to the linear function favored by the biochemical networks. Fits for assortativity favor a linear function for random reaction networks, whereas the assortativity of the biochemical networks was found to not scale with size (Supplementary Data S1). That is, the topology of random reaction networks scales with network size in a manner that is entirely distinct from that of real biochemistry, even though the two network ensembles (random and biological) share the same global set of chemical reactions. In addition to these qualitative differences in scaling behavior we also observed statistically significant quantitative differences in the random chemical networks: scaling relationships for randomly sampled biochemical networks do not overlap with real biological individuals in many cases. Topological measures in random chemical networks which scale according to power-law fits (y = y_0_ x^β^, β=β_ran_ for random reaction networks) include: average clustering coefficient (β_ran_=0.6401), average betweenness (β_ran_=-1.0595), average shortest path length (β_ran_=-0.0543), and number of edges (β_ran_=1.2459). Both biochemical diversity measures also scale according to power-law fits: number of enzyme classes (β_ran_=1.10156), and number of reactions (β_ran_=1.3590). Assortativity scales with a linear fit (y = mx + y0; m_ran_=-4.5255). The differences in scaling behavior indicate the real and random biochemical networks represent different universality classes. We conclude the organizational properties of random chemical networks, and in particular the existence of shared biochemistry across all organisms on Earth, cannot alone explain the scaling laws observed for reach biochemical networks.

**Figure 4:**
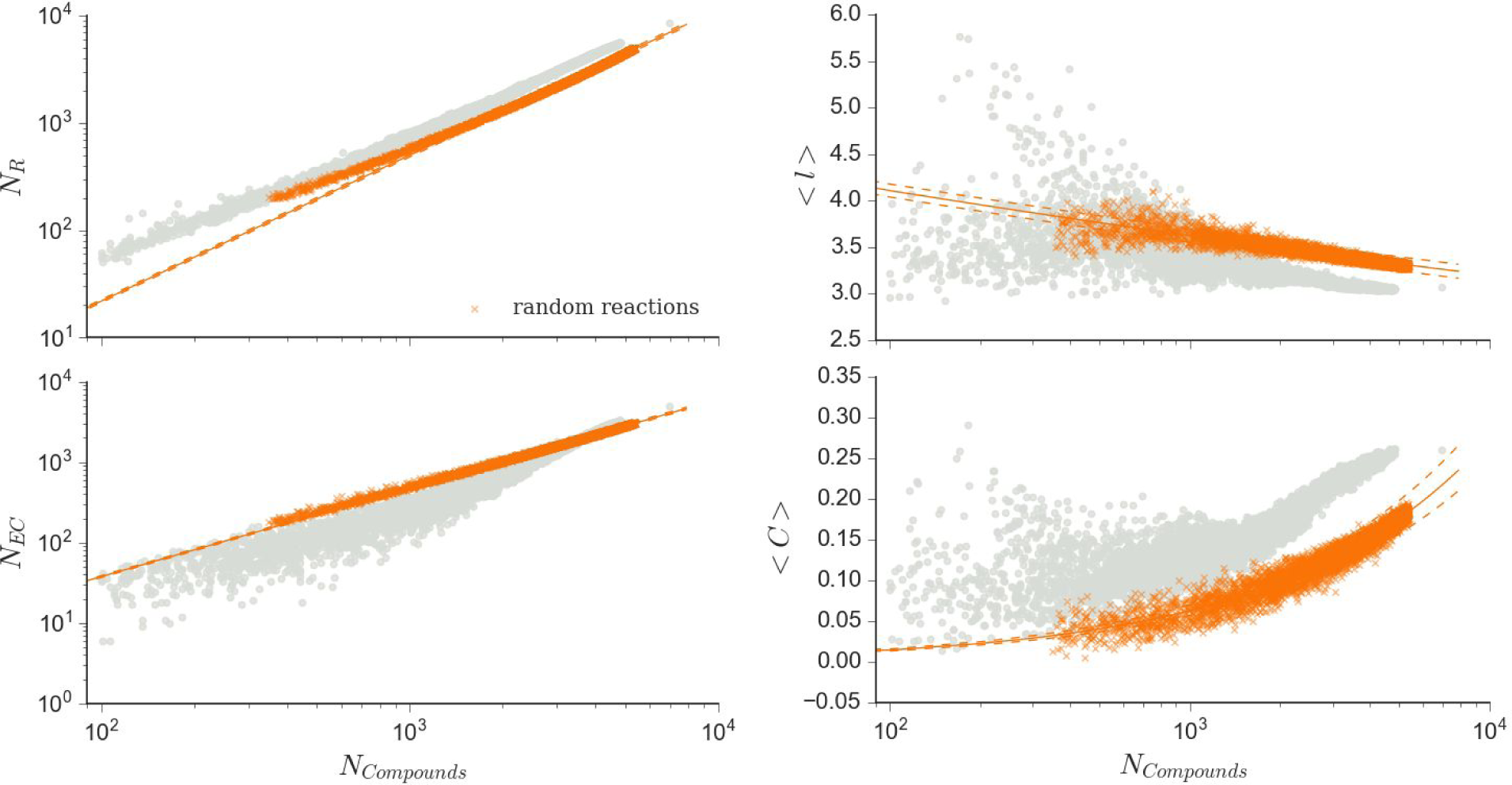
Scaling laws distinguish biochemical networks from random networks across levels of organization. Catalytic diversity scaling (left column) and topological scaling (right column) for merged networks composed of randomly sampled reactions cataloged in KEGG (orange, right column). Measure and fit descriptions match those described in Fig. 3. For reference, all real biochemical network data from Fig. 3 is shown in light grey. Additional measures are shown in Supplementary Fig. S5, and scaling for bipartite networks shown in Supplementary Fig. S6.

### Scaling-laws represent shared constraints re-emerging across levels

Our results establish biochemistry exhibits universal scaling behavior across levels of organization not explainable by shared biochemistry across life alone. A natural next question is whether ecosystems inherit their properties from individuals, or whether they instead exhibit similar structure due to similar constraints re-emerging at different levels. To address this, we next sought to determine whether or not scaling behavior for individuals is statistically distinguishable from ecosystems. We assumed as a null hypothesis scaling relationships consistent across levels of organization and performed a permutation test^57^, using the scaling coefficient as the test statistic (see Supplementary methods on *Fitting network measure scaling and permutation tests*). We find scaling relationships are not distinguishable for individuals and ecosystems when analyzing average node betweenness and average shortest path length (Fig. 5 and Supplementary Table 3). However, scaling coefficients are distinguishable for number of reactions, number of edges, number of enzyme classes, and mean clustering coefficient, with p-values < 10^−5^ in most cases. Confidence intervals on scaling coefficients for ecosystem topology are narrower than for individuals, indicating ecosystem scaling is more tightly constrained. Although biochemical networks for individuals and ecosystems share similar scaling behavior, they are not drawn from the same distributions; allowing the possibility shared constraints operate at each level separately.

**Figure 5:**
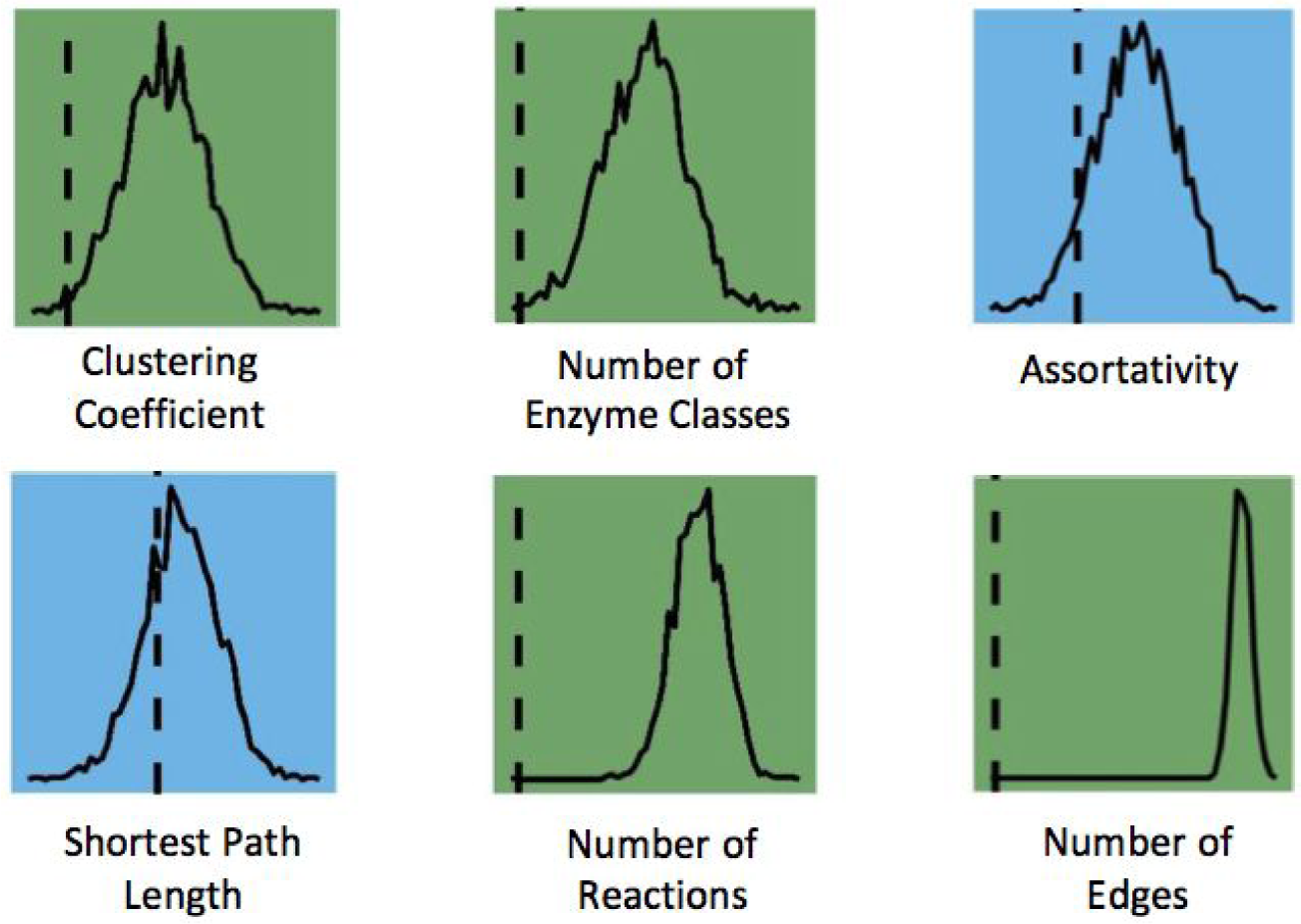
Scaling laws for individuals and ecosystems are statistically distinguishable for some network and catalytic diversity measures. Shown are the results of a permutation test to determine whether properties of biochemical networks constructed from individual genomes scale differently than those constructed from metagenomes (ecosystems). For each network measure the test statistic is shown as a vertical dashed line, while the null distribution is shown as a solid line (see Methods on *Fitting network measure scaling and permutation tests* for more details). Blue squares indicate scaling behavior is indistinguishable between levels of organization, while green squares show measures which can distinguish scaling of individuals from that of ecosystems.

We next generated simulated ecosystem-level networks by merging randomly sampled genome networks from each domain individually and from all three domains together (see Supplementary Materials and Methods for details on network construction). This allows us to determine how scaling behavior could be the same or different for an archaeasphere (archaea alone), bacteriasphere (bacteria alone), eukaryasphere (eukarya alone), or artificial ecosystems (all three domains) (Fig. 6). We find the functional form of scaling relationships are the same for real ecosystems and randomly merged organismal networks (hereafter called random genome networks). This is somewhat surprising given it is not in general true randomly selected subnetworks of a network have the same structure as the original network ^58^ (e.g., individuals as subnetworks of ecosystems do not necessarily have to share the same structure).

**Figure 6:**
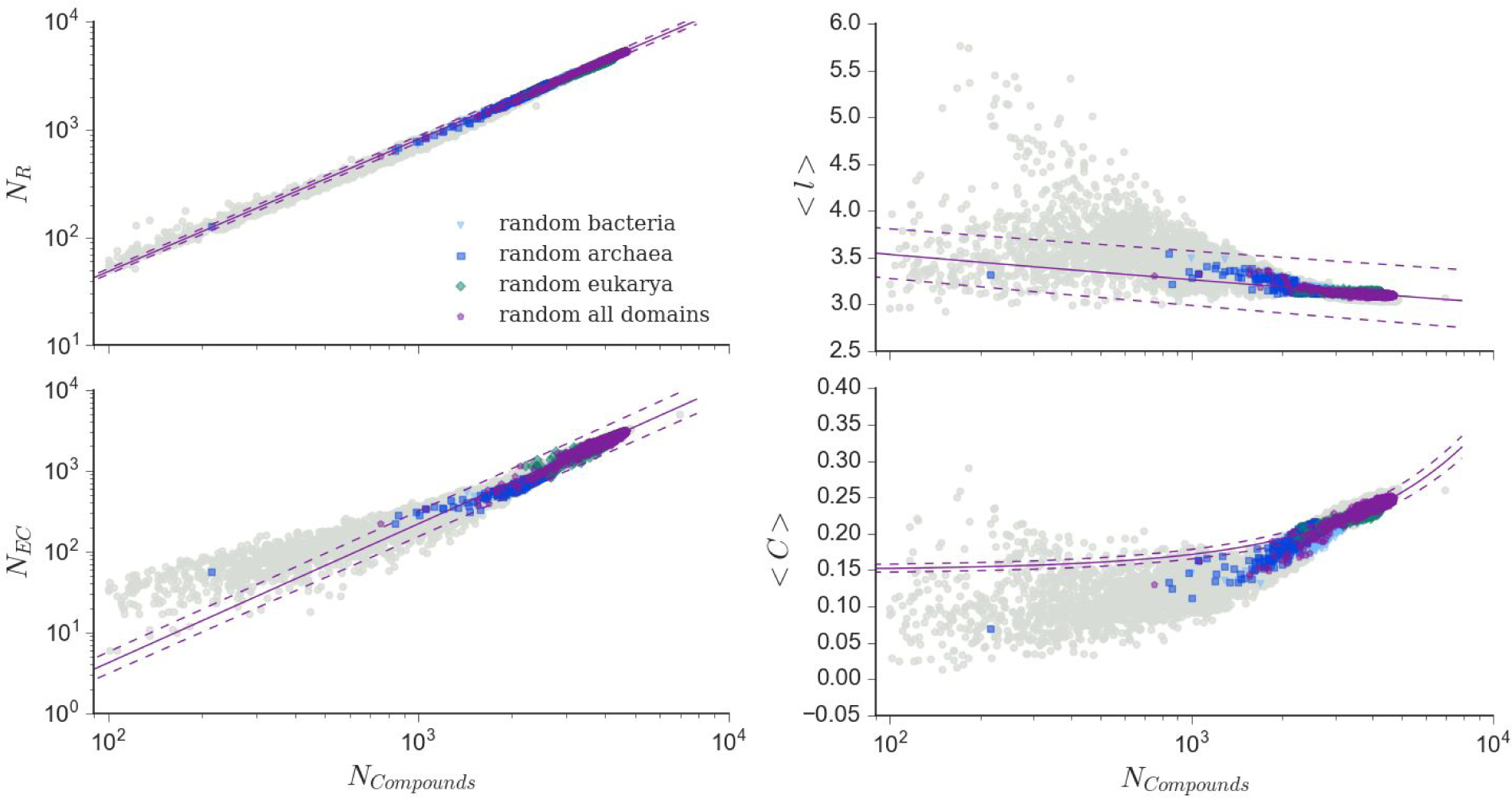
Scaling laws for random genome networks generated by merging biochemical networks of randomly sampled individuals from the three domains of life. Shown are catalytic diversity scaling (left column) and topological scaling (right column). Measure and fit descriptions match those described in Fig. 3. Merged networks composed of individuals include bacteria only (light blue), archaea only (dark blue), eukarya only (blue-green), and all domains combined (purple). For reference, all real biological networks from Fig. 3 are shown in light grey. Additional measures are shown in Supplementary Fig. S5, and scaling for bipartite networks shown in Supplementary Fig. S6. We find ecosystem-level biochemical networks and random genome networks scale with the same fit, but are statistically distinguishable for most measures.

One explanation for shared structure of random genome networks and real ecosystems is the common chemistry shared across all life. But we have already ruled this out as the explanation for the observed scaling in real ecosystems in the previous section. Combining these results with those of the previous section therefore indicates the existence of individuals, as specific partitions of the biosphere-level network shaped by selection, are sufficient for networks of all sizes (from small individuals to large ecosystems) to exhibit the scaling behavior observed in real living systems. Whether biological individuals are *necessary*, as opposed to being simply *sufficient ent* for recovering the observed scaling remains an open question.

We find scaling exponents and coefficients are similar for real ecosystems and random genome networks. However, we also checked whether they are statistically distinguishable, using the same permutation tests as before, and find they are for most measures (see Supplementary Table S2). Random genome networks and real ecosystems exhibit exponents distinguishing their scaling coefficients for most topological measures and for number of enzymes with p-values < 10^−5^. Scaling of betweenness is indistinguishable between the two datasets. These results indicate random genome networks differ from real ecosystems in many of the same ways individuals do. However, scaling of assortativity does distinguish random genome networks from real ecosystems, whereas it does not distinguish individuals from ecosystems. Taken on the whole, these results suggest scaling behavior for ecosystems arises due to selection on network architecture existing at the level of ecosystems, and is not solely an emergent property due to merging individual-level networks.

### Network structure predicts evolutionary domain

Any general organizing principles in biology must be consistent with the variation responsible for the diversity of life we are already familiar with. Since the three domains of life represent the most significant evolutionary division in the history of life ^59^, we therefore tested whether or not network structure can distinguish individuals sampled from the three domains (see Methods on *Predicting evolutionary domain from topology*). To approach this question, we first investigated compounds shared across all domains to determine which compounds are distinct to each domain and which are universal to all three. We identified the contributions of each domain to the biosphere as a whole by comparing compounds at the biosphere-level to those across the networks of individuals, identified by evolutionary domain. We do so by identifying which compounds are unique to each domain and which are shared across all three domains, determined from annotated data in the 22,559 genomes in our dataset. At the biosphere-level, 0.44% of compounds are unique to archaea, 3.14% are unique to bacteria, and 17.08% are unique to eukarya, reaffirming each domain represents significantly different metabolic strategies and genetic architectures, as is well established by earlier work (Hug et al. 2016). However, it is also well established all life on Earth shares a common set of core-biochemistry ^60^: a higher percentage of compounds, constituting 37.23% of the biosphere-level network, are shared across all three domains in our dataset (Fig. 7), including hubs such as ATP, and H_2_O as mentioned previously. Since many more chemical compounds are shared across all three domains than are unique to each, one might a priori expect the organization of these compounds into biochemical networks to *not* be predictive of domain.

**Figure 7:**
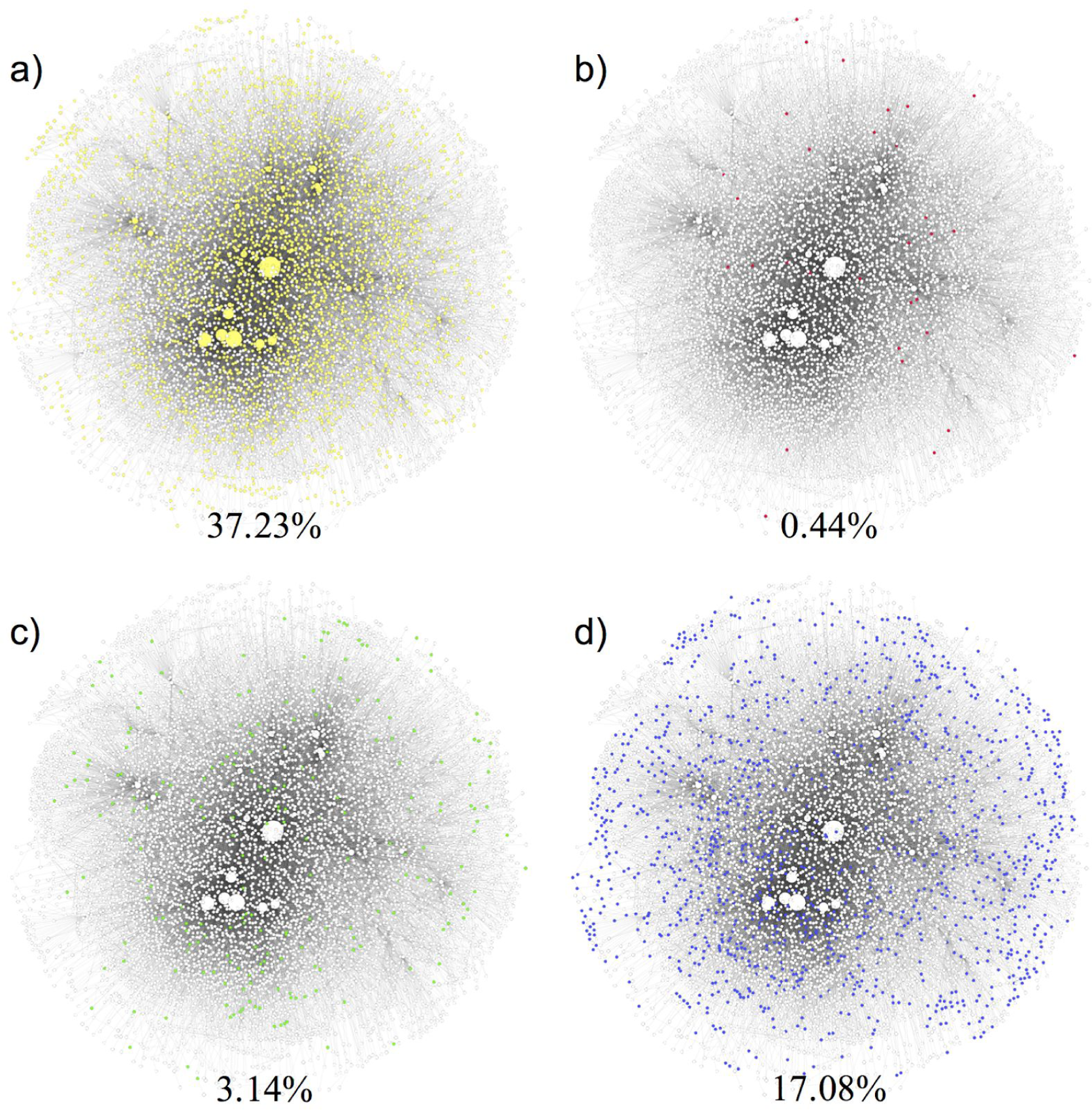
A biosphere-level chemical reaction network, constructed from the union of all 22,559 genomic networks in our study. Each panel shows the same biosphere-level network where nodes are white and edges are grey. Node size indicates its degree within the network. Colors indicate biochemical compounds used in (a) all three domains of life (yellow), (b) in archaea only (pink), (c) in eukarya only (green) and (d) in bacteria only (blue). Although many more chemical compounds are shared across all three domains than are unique to each, the organization of these compounds into biochemical networks is distinct for each domain (see Fig. 8).

We find the opposite to be true: despite a large fraction of shared biochemical compounds, the organization of those compounds into networks is distinct for each domain. We find in most cases average topology normalized to size can reliably predict evolutionary domain (Fig. 8). In many cases prediction accuracy is > 80%, when only a single topological measure is used. By contrast, topology or size alone provides significantly less accurate predictions. This demonstrates biochemical network structure can be predictive of the taxonomic diversity of individuals. Combined with our other results this suggests the same biochemical network properties (topology and catalytic diversity) driving regularity *across* levels of organization can also be predictive of major evolutionary divisions *within* a given level, providing evidence the global organization of biochemistry is indeed consistent with a signature of selection.

**Figure 8:**
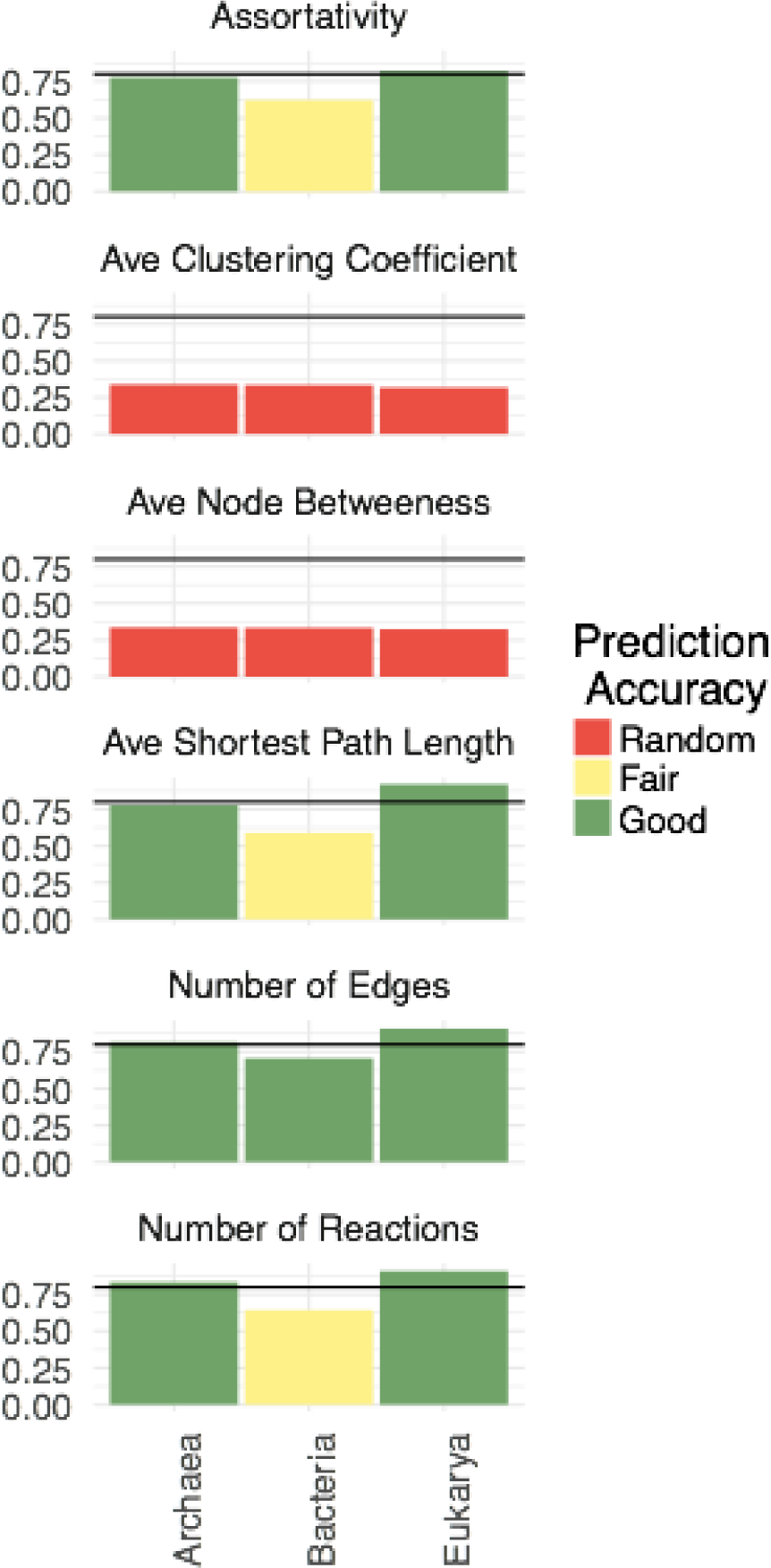
Catalytic diversity and biochemical network topology can predict evolutionary domain. Shown is the estimated prediction accuracy (y-axes) for each measure and each domain. The colors of each bar indicate prediction accuracy of a given measure for a particular domain: red is comparable to random guessing (y<=33% accuracy); yellow is better than random but not completely predictive (33%<y<=67%); green is predictive of domain (67%<y). The horizontal line indicates 80% prediction accuracy.

## Discussion

Our analyses reveal biochemical networks display common scaling laws governing their topology and biochemical diversity, which are independent of the level of organization they are sampled from and not explained by the structure of random chemical networks. We were also able to confirm the same regularities occurring across levels of organization within the the biosphere can be predictive of evolutionarily divisions within a level, using the three domains as an exemplar. Collectively, our results indicate a deeper level of organization in biochemical networks than what is understood so far, providing a new framework for understanding the planetary-scale organization of biochemistry and how selection structures nested hierarchies.

A key implication of our analysis is the importance of hierarchical levels and selectable units in shaping the universal scaling laws observed across biochemical networks. Scaling laws often emerge in systems where universal mechanisms operate across different scales, yielding the same effective behavior independent the specific details of the system. It is in this sense scaling laws can uncover universal properties, motivating their widespread use in physics and increasing application to biology^42,46,48,61–64^. Here we have shown the relevant scaling parameter for biochemical organization is the number of biochemical compounds (in a network representation this is the size of the network). Individuals, ecosystems and the biosphere obey much the same scaling behavior for biochemical network structure, indicating the same universal mechanisms could operate across all three levels of organization. In physics, this kind of universality usually implies there is no preferred scale or basic unit. However, in the biological example uncovered here, the presence of specific scaling relations observed in real biochemical networks can be explained by biological individuals as a basic ‘units’. That is, somewhat paradoxically individuals seem to be sufficient for biology to exhibit the observed scaling behavior across levels. Random chemical networks, even if they share the same global biochemistry as our biosphere, exhibit different scaling behavior, perhaps reflective of differences in the universality classes of biochemical networks and random chemical networks.

Future work should explore the connections between the scaling relationships reported here and other work characterizing scaling behavior across living processes. For example, our results indicate ecosystems are more tightly constrained than individuals, better displaying the regularities of biochemical network architecture. However, projecting ecosystem-level scaling to the biosphere as a whole does not recover the observed network properties for the biosphere-level network. Recently, scaling laws describing microbial diversity were used to predict Earth’s global microbial diversity, and in particular to highlight how much diversity remains undiscovered ^44^. It could be an analogous case here, where scaling relations predict missing enzymatic diversity in the biosphere. Furthermore, one area of intensive investigation is allometric scaling relations^44,61,64^, including how shifts in metabolic scaling could be indicative of major transitions in evolutionary hierarchies ^42^. Allometric scaling laws are derived by viewing living systems as localized physical *objects* with energy and power constraints. Here, scaling emerges due to an orthogonal view of living systems as distributed *processes* transforming matter within the space of chemical reactions. The connection between these different aspects of scaling in living organization remain to be elucidated.

A final implication of our work are the consequences for our understanding of the origin of life, before the emergence of species. The existence of common network structure across all scales and levels of biochemical organization suggests a logic to the planetary-scale organization of biochemistry^65^, which - if truly universal - would have been operative at the origin of life. An important test of this hypothesis will be to determine whether the same global network structure, characterized by the same scaling laws, described Earth’s biosphere throughout its evolutionary history. If this is indeed the case, the emergence of individuals (as selectable units) would have played an important role in mediating a transition in the organization of Earth’s chemical reaction networks. Even if we could assume the same planetary-scale chemistry for a lifeless world, we should expect to see dramatically different scaling for a hierarchically organized biosphere of nested evolutionary units ^66,67^. An important question for future work is identifying the planetary-drivers of Earth’s biosphere-level biochemical network structure and how this has structured living systems across nested levels over geological timescales. This will require characterizing the organization of planetary-scale biochemistry, as developed here, within the broader context of studying a planet’s geologic and atmospheric evolution. It remains an open question as to what will ultimately explain the universal structure of Earth’s biochemical networks, or whether we should expect all life to exhibit similar scaling behavior, even on other worlds.

## Acknowledgements

We thank the Emergence@ASU team for feedback through various stages of this work and are grateful for support from the National Aeronautics and Space Administration through grant NNX15AL24G S02, which made this research possible.

## Methods and Materials

### Obtaining genomic and metagenomic information

#### Genomes (PATRIC)

Archaea and bacteria genomic datasets were obtained from PATRIC^30^. Enzyme commision (EC) numbers were obtained from “ec_number” column in the pathway data of each taxon. Eukarya genomic datasets were obtained from the Joint Genome Institute’s (JGI) integrated microbial genomes database and comparative analysis system (IMG/M)^31^. All eukarya data used in this study was sequenced at JGI. All EC numbers used to construct eukarya biochemical networks were obtained from the list of total enzymes associated with each eukaryote. EC numbers were used in conjunction with KEGG enzyme and reaction data in order to build biochemical networks for each taxon.

#### Metagenomes (JGI)

Metagenomic data was obtained from JGI IMG/M(Markowitz et al. 2012). All metagenomic data used in this study was sequenced at JGI. All EC numbers used to construct metagenomic biochemical networks were obtained from the list of total enzymes associated with each metagenome. These EC numbers were used in conjunction with KEGG enzyme and reaction data in order to build biochemical networks for each metagenome.

#### Biosphere

To create the biosphere network, we included all 8,658 enzymatically catalyzed reactions in KEGG.

### Network Construction

In this study, we construct three different types of biochemical reaction networks: biological networks, random genome networks and random reaction networks. These biochemical reaction networks consist of chemical compounds that are involved in biochemical reactions: two chemical compounds are connected to each other when one is a reactant and the other is a product of the same biochemical reaction. The process to encode a biochemical reaction as the network representation can be described with the diagram below as follows:

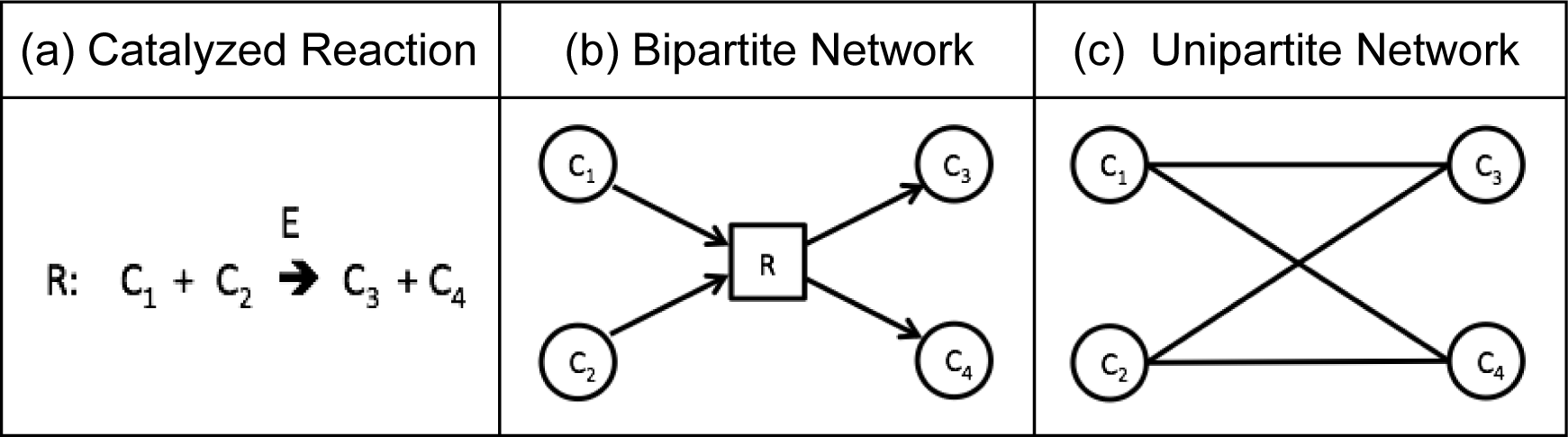

(a) Suppose that a chemical reaction R catalyzed by an enzyme E is given, which transforms chemical compounds C1 and C2 to C3 and C4. (b) The reaction, R, can be described in a reaction diagram, or a directed bipartite network representation, where the reactants C1 and C2 are connected to the reaction node and the products C3 and C4 are connected as products from the same reaction. In principle, this biochemical reaction, R, can happen in opposite direction depending on the environment. Therefore, in bipartite network representation, the edges connecting chemical compounds and the reactions are considered as bidirected, which is equivalent to undirected for our analysis. (c) The unipartite network representation of the reaction, R, shows how the reaction information is embedded in the network. In the unipartite network representation, nodes are substrates and a reactant is connected directly to a product if they are connected to the same reaction in the corresponding reaction diagram.

Regardless of the types of networks, all chemical network representations in this paper follow the same methods. Therefore, the distinctions between different types of biochemical reaction networks come from how we select reactions to be included in each network, which is described below. Note that all edges in the networks in this paper are represented as undirected and unweighted since our interests lie on the presence or absence of particular reactions in given networks and, in principle, all biochemical reactions can happen in both directions depending on the environment.

#### Biological Networks

For each biological network, we include all catalyzed biochemical reactions annotated in each genome or metagenome. More specifically, we consider three different levels of organization: individual organisms, ecosystems and the biosphere. For the construction of individual networks, we utilize the genome data of 21,637 bacterial taxa and 845 archaeal taxa from the Pathosystems Resource Integration Center (PATRIC)^30^, as well as 77 eukaryotic taxa from the Joint Genome Institute (JGI)^31^. From this data, we obtain the set of classes of enzymes for each genome. All reactions catalyzed by this set of enzymes and present in the Kyoto Encyclopedia of Genes and Genomes (KEGG)^32^ database are included in the network representation of the corresponding genome.

Similarly, for the network representation of each of the 5587 ecosystems from JGI, we include all reactions catalyzed by the ecosystem’s coded enzymes, provided they are catalogued in the KEGG dataset. Finally, for the biosphere network, we include all 8,658 enzymatically catalyzed reactions in KEGG.

#### Random Genome Networks

To construct a random genome network, we sample individual networks uniformly at random from the set of all individual organisms in our data set and merge them into one random genome network. When a set of multiple individual networks are merged, every node and edge present in any individual network are added to the resulting network with equal weight regardless of how many individual networks include them. We built 4 types of random genome networks with individual networks sampled from only archaea, only bacteria, only eukarya and from integration of all the three domains. In total, we generated 2,000 random genome networks from 730 individual archaea networks, 2,000 from 21,213 individual bacteria networks, 770 from 77 individual eukarya networks, and 4770 from merging all individual networks in the three domains.

#### Random Reaction Networks

In this paper, random reaction networks are generated by merging randomly sampled reactions from all biochemical reactions from KEGG data regardless of whether a known enzyme is cataloged for the reaction. We note 31.46% of chemical compounds in the biosphere network are not included in the genomic data in our study, therefore our construction uniformly sampling the entire KEGG database, the random reaction networks can include enzymatically catalyzed reactions not included in our genomic data. Nonetheless our sampling procedure is biased to generate networks with similar biochemistry to that of the genomic networks, due to reasons explained in the main text (compounds common to all three domains tend to be highly connected (participate in many reactions) such that a uniform sampling procedure yields random networks biased to include the most common compounds used by life). Most biological networks for real individual organisms and ecosystems contain 200 - 5000 reactions. Hence, to build similar size of random reaction networks to real individual organisms and ecosystems, we selected the total number of reactions in each network from the range between 200 and 5000, sampling for each size the appropriate number of reactions from KEGG data uniformly and at random. Merging these into networks, we constructed 5,000 random reaction networks in total.

#### Topological Measures

To characterize the topology of biochemical networks, we utilized some of the most frequently used topological measures. The detailed descriptions about these topological measures can be found in^37^. Below, we briefly review these measures and related terms. For computing each measure, we used functions provided by Python software package, NetworkX^68^. The topological measures implemented in this paper include average degree, average clustering coefficient, average shortest path length, assortativity (degree correlation coefficient), and node betweenness.

We calculate all network measures on the largest connected component (LCC) of each network, for the following reasons: 1. Several network measures only make sense to calculate on connected components (e.g. average shortest path), focusing on the LCC therefore permits all network measures implemented in our study to be calculated for all networks; 2. The largest connected components have the vast majority of nodes (>90%) for the vast majority of networks in each dataset (the only exception is the random reaction networks, of which only ∼76% have a largest connected component with at least 90% of a network’s nodes). See **Table S1** and **Fig. S1** for distribution of sizes of the LCC by dataset.

#### Degree

The degree of a node *i*, *ki* is the total number of connections between *I* and rest of the network. The average degree < *k* > in this paper is average of *ki* for all nodes in LCC of a given network.

#### Clustering coefficient

The local clustering coefficient for a node *i*, *C*_*i*_ measures the local density of edges in a network by considering the number of connected pairs of neighbors of defined as,

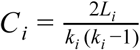

where *k*_*i*_ is the degree of node *I* and *L*_*i*_ is the number of connections between neighbors of *i*. The large values of *C*_*i*_ indicates the highly interconnected neighborhood of *i*. *C*_*i*_ is measured by using a Networkx method **clustering(..)** and we computed < *C* >, the average of *C*_*i*_, over all nodes in LCC of each network.

#### Shortest path length

The shortest path length,*l*_*ij*_ between a given pair of two nodes *I* and *j* is defined as the minimum number of edges connecting the two nodes in a given networ*l* ^*ij*^ is measured by using a Networkx method **shortest_path_length(..)**. We calculated the average shortest path length,.< *l* > by averaging *l*_*ij*_ for every pair of nodes in LCC of a given network

#### Assortativity (degree correlation coeffiCient)

Assortativity measures the tendency of two nodes with similar properties to be connected in a given network. The assortativity coefficient proposed by Newman^69^ is formulated as follows:

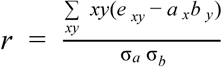

Where *e*_*xy*_ is defined as the fraction of edges between a node with value *x* and one with value *y* for a given node attribute, and *a*_*x*_ and *b*_*y*_ are the fraction of edges coming into and going out from nodes of value *x* and *y* respectively. σ_*a*_ and σ_*b*_ are the standard deviations of the distributions of *a*_*x*_ an *b*_*y*_. When the considered attribute of nodes is their degree, the assortativity becomes the degree correlation coefficient, the correlation between the degrees of nodes on either side of an edge. Hence, for undirected networks in our study, *ax* = *b*_*y*_ and σ_*a*_ σ*b* = σ^2^. If *r* < 0, the network is assortative,i.e. nodes with similar degree tend to be connected to each other. If *r* > 0, the network is disassortative, i.e. nodes in it tend to be paired to other nodes with different degrees. For an arbitrary network, −1 ≤ *r* ≤ 1. To measure the assortativity *r*, we used a Networkx method **degree_assortativity_coefficient(..)**.

#### Betweenness

Betweenness centrality of a node, *B*_*i*_ is defined as ^70^,

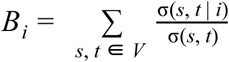

Where *V* is the set of all nodes in a network, and σ(*s*, *t*) and σ(*s*, *t* |*i*) denote the number of all shortest paths from *s* to *t*, and the number of the shortest paths through a given node *i*, respectively. Replacing σ(*s*, *t* |*i*) with σ(*s*, *t* |*e*) for an edge *e*, one can also formulate the edge betweenness. *Bi* measures degree of importance of for the interactions between subsets of a given network. To compute *Bi*, Networkx methods **betweenness_centrality(..)** is implemented and < *B* > over every is average of *B*_*i*_ node in LCC of a given network

### Fitting network measure scaling and permutation tests

For each network measure, a scaling relationship was fit as a function of the size of the largest connected component (LCC) of the network. For each measure, three different models were tested, a power law of the form y = y_0_ x^β^, a linear relationship of the form y = β_x_ + y_0_, and a quadratic function of the form y = β_1_x + β_2_x^2^ + y0, for both the assortativity measures, the preferred fit was also compared to a constant y = β. The preferred model was chosen as the one which minimized cross validation errors, according to 10-fold cross validation, across the entire data set.

Once a model was chosen, a simulated permutation test was performed to determine whether the scaling relationship for a given attribute was the same for ecosystems and individuals or if it was distinct ^57^. We took as the null hypothesis the scaling relationship across different levels of organization are constant, and used the fitted scaling parameters (for individuals and ecosystems) as the test statistic. We used fitted 1,000,000 resamples of the complete dataset to estimate the likelihood of the fit for individuals (or ecosystems) to have been drawn randomly from the complete dataset. We performed this test for both the ecosystem and individuals, if there was a difference in the estimated likelihoods we took the greater of the two. These likelihoods are the (two-sided) p values reported in **Table S2**. The same procedure was followed to determine the distinguishability of ecosystem networks with the randomized controls (random genome networks, and random reaction networks). Random reaction networks were distinguishable from ecosystems networks for all measures, with p values = 10^−6^.

To estimate the true scaling parameters, and 95% confidence intervals a bootstrap sample of 100,000 was used for each network attribute ^57^. If the permutation test allowed us to reject the hypothesis of a constant scaling relationship across individuals and ecosystems to a confidence greater than 0.01, the scaling parameters were estimated separately for the individuals and ecosystems, otherwise the complete dataset was fit. The scaling parameters (and confidence intervals) for distinct domains were also estimated using a bootstrap of 100,000 samples.

For scaling fits and confidence intervals see **Table S2**.

### Predicting evolutionary domain from topology

To demonstrate topological features of genomes from different domains are distinct, multinomial regression was used. specifically, we implemented models where the domain of the network was the response class and a single topological feature, normalized by the size of the largest connected component (LCC) of the network was the dependent variable. We found topological features of networks alone were often not predictive of the domain but the ratio of the topological properties to the size of the network provided a more accurate prediction. Prior to the regression these normalized topological measures were scaled and centered ^57^. The regression was implemented in base R using the **glm(..)**, function. In order to control for over fitting the training data was composed of an equal number of samples from each domain. In particular only 35 networks of each domain were sampled and the model was tested on the remaining data. This process was repeated 100 times and the average model error is reported in the **main text figure 8**.

